# Early visual experience redistributes *cFos*-tagged neurons toward the cortex while preserving presynaptic spatial stereotypy

**DOI:** 10.64898/2026.11.08.765223

**Authors:** Oliver J. L’Esperance, Daniel Ryskamp Rijsketić, Nicholas Pritchett, Tristan Meier, Ethan Hunt, Boris D. Heifets, Jaichandar Subramanian

## Abstract

The macroscale functional organization of the brain is spatially stereotyped across individuals, but whether this extends to finer spatial scales is not known. Furthermore, how experience shapes this organization across the brain remains unclear. To address this, we developed *cFos* promoter-driven reporter mouse lines that permanently label either the cell bodies or presynaptic terminals of neurons during defined temporal windows. Using serial two-photon tomography and voxel-wise statistics, we mapped brain-wide distributions of *cFos*-tagged somata or their presynaptic terminals during the first week of visual experience or its deprivation. We found that midbrain and subcortical regions show greater stereotypy for *cFos*-tagged neurons and their presynaptic terminals compared to cortical regions. This fine-scale spatial stereotypy remained largely preserved despite visual deprivation, which shifted the relative recruitment of *cFos*-tagged neurons from cortical toward the highly stereotyped midbrain and subcortical regions. Our findings identify heterogeneous, spatially constrained functional organization across the brain that is conserved across subjects yet accommodates experience-dependent reorganization of neuronal recruitment.

## Introduction

Experience shapes brain development by altering how neural activity is distributed across interconnected brain regions ^1–6^. Whole-brain atlases have been created to identify cell types ^7–9^, connectivity ^10–14^, and brain-wide activity and metabolism ^15–19^, enabling understanding of how experience ^14,16,20^, development ^21–23^, or disease ^24,25^ modify the entire brain beyond individual neurons or local circuits. Whole-brain functional mapping in the mammalian brain is typically performed using functional magnetic resonance imaging ^10,13,25,26^ or immediate early gene mapping, which provides cellular resolution characterization of diverse physiological conditions ^14,16,20,24,27–37^. Primarily, these studies identify brain regions recruited during specific experiences by measuring changes in the density of neurons exhibiting *Fos* promoter activity. Typically, this is performed by assigning cells to predefined brain regions to compare regional cell density. To avoid missing meaningful biological changes due to prespecified regional constraints, UNRAVEL ^28^ was developed to register brains to a common atlas space and perform unbiased voxel-wise statistics that control for the false discovery rate (FDR) ^38^ across millions of voxels compared across the brain. Together, *cFos*-tagging, considered a proxy for salient activity ^27,39^, whole-brain imaging ^16,40^, and voxel-wise, quantitative topographic mapping ^28,41^ provide a powerful tool to explore experience-dependent functional reorganization across the brain.

The recruitment of different brain regions by experience could be influenced by their organization. Brain regions differ in their cytoarchitecture, cell type composition, and spatial organization ^7,8,42,43^. Cortical regions possess laminar organization ^44,45^, whereas many subcortical regions are organized as nuclei and subnuclei with complex spatial boundaries ^43,46^. Even within a single anatomical region, neurons with different molecular identities or projection targets can have non-uniform spatial arrangements ^46–49^. These organizational features could constrain where active neurons are recruited within a region. Therefore, the precise spatial location of experience-recruited neurons could become stereotypical between subjects, as evidenced by the common topography of activity tagged neurons in the amygdala ^48,50–52^, hippocampus ^27,53,54^, and olfactory bulb ^55–57^. However, the extent to which spatial stereotypy (between-subject variance) varies across the brain at the micrometer scale, whether it is shaped by experience, and whether intrinsic differences in stereotypy between brain regions predict their experience-dependent reorganization of activity-tagged neurons remain unknown.

Previous studies reporting changes in the average density of activity-tagged neurons in specific brain regions have captured one component of experience-dependent circuit reorganization. However, neurons also influence downstream circuits through their axons and presynaptic terminals. Therefore, the spatial distribution, stereotypy, and experience-dependent reorganization of presynaptic terminals of activity-tagged neurons may differ substantially from their soma, since these projections are structured to target specific nuclei or layers across the brain. To date, brain-wide organization of presynaptic terminals of activity-tagged neurons has not been imaged, and consequently, the extent to which their distribution and stereotypy are modulated by experience is not known.

To address these gaps, we developed an approach using UNRAVEL ^28^ to map the brain-wide organization of neurons exhibiting *cFos* promoter activity (via *cFos-*dependent tdTomato production) and their presynaptic terminals (via *cFos-*dependent synaptophysin-tdTomato production), to quantify their spatial stereotypy across subjects, and to determine how salient changes to early visual experience reorganize this cellular and presynaptic organization.

## Materials and methods

### Mice

All animal procedures are approved by the University of Kansas Institute of Animal Use and Care Committee and meet the NIH guidelines for the use and care of vertebrate animals. *cFos-tTA* mice (*B6.Cg-Tg[Fos-tTA,Fos-EGFP*]1Mmay/J*, RRID: IMSR_JAX:018306)^39^, *TetO-Cre* mice (Cre-dependent tdTomato; *B6.Cg-Tg[tetO-cre]1Jaw/J*, RRID: IMSR_JAX:006234)^58^, *Ai14* mice (*B6.Cg-GT[ROSA]26SOR^tm14[CAG-tdTomato]Hze^*/J, RRID: IMSR_JAX:007914)^59^, and *Ai34* mice (Cre-dependent synaptophysin-tdTomato; *B6;129S*-*Gt[ROSA]26SOR^tm34.1[CAG-Syp/TdTomato]Hze^/J*, RRID: IMSR_JAX:012570)^60^ were originally obtained from the The Jackson Laboratory, an NIH-funded strain repository. *cFos-tTA* heterozygotes were crossed with *Ai14* or *Ai34* homozygotes to generate double heterozygotes, which were then backcrossed to their respective *Ai14* or *Ai34* line to generate *cFos-tTA^+/-^Ai14^+/+^* or *cFos-tTA^+/-^Ai34^+/+^* mice. These mouse lines were maintained by continued backcrossing to *Ai14* or *Ai34* homozygotes. To generate triple transgenic experimental line mice (*cFos-tTA^+/-^TetO-Cre*^+/-^*Ai14^+/-^*or *cFos-tTA^+/-^TetO-Cre^+/-^Ai34^+/-^*), *cFos-tTA^+/-^Ai14^+/+^* or *cFos-tTA^+/-^Ai34^+/+^* mice were crossed with *TetO-Cre* heterozygotes. All breeder and experimental mice were genotyped to confirm the presence of relevant genes. A maximum of five mice were housed in a standard cage and maintained on either a 12h light/12h dark cycle or in 24h light deprivation. For light deprivation, mice were placed in a ventilated and lightproof cabinet inside of a dark room. To prevent any light exposure, infrared goggles with a low-intensity 850nm wavelength emission source were always used while maintaining mice in dark housing.

### Experimental timelines

For *Ai34* line triple transgenic signal and noise characterization, mice were maintained on 40mg/kg doxycycline in mouse chow (Teklad doxycycline diet TD.120240 [40 ppm Doxy Diet 2018, B]) until either postnatal day 2 (P2; dox^-^) or until P30 (dox^+^). Both groups were transcardially perfused on P30. For other experiments, mice were maintained in a 12h light/12h dark cycle from birth to P8 followed by visual deprivation from P9-14 to prevent diffuse light stimulation through the eyelids. On P15, pups were visually verified for eye opening with infrared goggles in the dark, and mice were either maintained in a 12h light/12h dark cycle (Light group) or 24h darkness (Dark group) until P28 transcardial perfusions. For these Light vs. Dark serial two-photon tomography experiments, mice were maintained on doxycycline until P10. Mice were fed doxycycline-free mouse chow from P10 to P14 to allow for doxycycline clearance prior to eye opening. Doxycycline-free chow was continued until P21, then replaced with 40mg/kg doxycycline chow to halt further *cFos* promoter-dependent fluorescent labeling until P28 transcardial perfusions and brain collections. For parvalbumin experiments, a separate set of mice were housed in exactly the same Light or Dark conditions until P28 transcardial perfusions.

### Tissue preparation

Mice were deeply anesthetized by intraperitoneal injection of 2% avertin in phosphate-buffered saline (PBS), pH 7.4, and transcardially perfused with cold PBS followed by 4% paraformaldehyde. The brains were extracted and post-fixed in 4% PFA overnight at 4°C, followed by storage in PBS. The brains were embedded in 4% oxidized agarose as previously described^20^ to limit artifacts during sectioning. For P2-30 signal/noise characterization, parvalbumin immunohistochemistry, and *cFos-tTA^-/-^TetO-Cre^+/-^*Light vs. Dark experiments, the brains were coronally sectioned at a thickness of 40µm with a vibratome (Leica VT1000 S). Serial two-photon tomography brains were coronally sectioned and imaged at 100μm intervals.

### Immunohistochemistry

Coronal brain sections were sequentially placed into 24-well plates submerged in 1x PBS and imaged on a Nikon Eclipse TE-2000U inverted microscope, using collected autofluorescence to identify individual slices containing brain regions/subregions of interest. 4-5 evenly spaced slices that spanned from the posterior midbrain to the anterior olfactory bulbs were selected per animal and fluorescently immunolabeled for parvalbumin-expressing cells. Slices were blocked for 2 hours at room temperature in 500μL per well of a PBS solution containing 10% NGS, 3% BSA, 1% Triton X-100, and 0.05% Tween-20 before proceeding to a 4°C overnight incubation with 5% NGS, 3% BSA, 0.1% Triton X-100, 0.05% Tween-20, and a mouse anti-parvalbumin (1:500, Sigma cat. # P3088). Slices were washed 5x with 0.05% Tween-20 in PBS and incubated for 2 hours at room temperature in PBS containing 5% NGS, 0.1% Triton X-100, 0.05% Tween-20, and an Alexa 647-conjugated goat anti-mouse antibody (1:2000, ThermoFisher cat. # A21236) to label parvalbumin. Slices were washed 5x in PBS prior to mounting on slides for imaging.

### Image acquisition

Immunostained brain slices and brain slices for P2-30 signal/noise experiments were imaged in widefield using an ImageXpress Pico automated imaging system (Molecular Devices, San Jose, CA) with a 10× objective (Leica HC PL FLUOTAR 10×/0.32). For the P2-P30 signal/noise experiments, twelve evenly spaced coronal slices spanning the posterior cortex to the olfactory bulb were used. For *Ai34* line *cFos-tTA^-/-^TetO-Cre^+/-^*Light vs. Dark experiments, eight evenly spaced coronal slices spanning from the posterior cortex to the anterior lateral geniculate nucleus were imaged in the same manner. For voxel-wise whole-brain analyses, the entire brains were coronally imaged at 100µm intervals using a TissueCyte 1000 serial two-photon tomography system^40^ (TissueVision, Newton, MA) with a 16x objective, yielding 1.2µm/pixel resolution with a 100µm z-step. Two-photon imaging with 1040nm excitation light, and emissions were filtered into >560nm, 500-560nm (tdTomato signal), and <500nm (blue autofluorescence) channels.

### Data and statistical analysis

For P2-30 signal/noise and *cFos-tTA^-/-^TetO-Cre^+/-^*Light vs. Dark experiments, brain section mapping into a common mouse brain atlas coordinate framework, *Ai34* line synaptophysin-tdTomato fluorescence collection, and brain region area measurements were performed using NeuroInfo software ^61^ (MBF Bioscience, Williston, VT). 12-bit images of brain sections were first mapped in 3D via both linear and nonlinear splining to the 2020 Allen CCFv3 atlas space ^43^ to allow for automated cell/synapse detection and area measurement by region. Fluorescence density was calculated in nine major anatomical subdivisions of the brain for P2-30 signal/noise experiments and compared across groups using a 2-way restricted maximum likelihood ANOVA (brain area [within-subject], genotype/dox combination [between subject], and their interaction) with post-hoc Tukey’s multiple comparisons tests across conditions within each brain area (9 families with 3 comparisons per family). Raw and z-scored fluorescence were calculated in five visual processing areas for *cFos-tTA^-/-^TetO-Cre^+/^*^-^Light vs. Dark experiments and compared between groups using a 2-way restricted maximum likelihood ANOVA with Geisser-Greenhouse’s epsilon correction matched by mouse (brain area [within-subject], lighting condition [between subject], and their interaction) with post-hoc Šídák’s multiple comparisons tests between conditions within each brain area (1 family, 5 comparisons each; all ns for raw and z-scored comparisons).

For *Ai14* line *cFos^-/-^TetO-Cre^+/-^*fluorescence vs. cell count vs. z-scored cell count correlation comparisons, pair-wise correlations (Pearson’s *r*) of either fluorescence, cell count, or z-scored cell count were calculated across 26 anatomical brain areas. For each brain area, the average Pearson’s *r* was calculated between it and every other brain region, then converted to z using Fisher’s r-to-z transformation. These regions’ combined average z-values were compared (when calculated from fluorescence vs. cell count vs. z-scored cell count, self-correlations omitted) using a one-way mixed-effects restricted maximum likelihood ANOVA with Tukey’s post-hoc multiple comparisons test (1 family, 3 comparisons). The coefficient of variance of values for each brain region across mice was likewise calculated, averaged across brain regions, and compared between groups (calculated from fluorescence vs. cell count vs. z-scored cell count) using a Friedman test (nonparametric CoV distributions), matched by brain region, with Dunn’s post-hoc multiple comparisons test (1 family, 3 comparisons).

*Ai14* and *Ai34* line tdTomato expression was mapped across serial 2-photon slices using UNRAVEL (UN-biased high-Resolution Analysis and Validation of Ensembles using Light sheet images; https://github.com/b-heifets/UNRAVEL) ^28^. Code was refactored to Python (documentation: b-heifets.github.io/UNRAVEL/index.html) and capabilities were extended to allow for serial 2-photon microscopy and cell centroid analyses for the *Ai14* line. Z-stacks with blue autofluorescence (excitation: 1040nm, emission: <500nm) were used for atlas registration. These stacks were resampled to a 50μm resolution, and tissue was segmented using Ilastik (v1.4.0; pixel classification with all features) to generate a mask for N4 bias field correction (ANTsPy 0.4.2). The resampled images were padded (15% of voxels on all sides), smoothed using a Gaussian filter (*σ* = 0.4) and registered with a serial 2-photon average template brain from the 2020 Allen Mouse Brain Common Coordinate Framework (CCFv3) ^43^, resampled to 30µm isotropic resolution. Atlas registration was performed using ANTsPy, first applying affine initialization (ants.affine_initializer(fixed_image=autofluorescence_image, moving_image=template, search_factor=1, radian_fraction=1, local_search_iterations=500)) followed by symmetric normalization (ants.registration(fixed=autofluorescence_image,moving=template_aligned_with_tissue, grad_step=0.1, syn_metric=’CC’, syn_sampling=2, reg_iterations=(100, 70, 50, 20))). Registration accuracy was visually assessed using FSLeyes (v0.31.0; FMRIB; available at https://zenodo.org/records/21164449) by overlaying the warped atlas on each registered image.

*Ai34* line images were warped to the 30µm atlas space using the transforms generated from the registration step. The images were smoothed using a 100μm kernel and mirrored to average the left and right hemispheres. To correct for background fluorescence and potential reporter leakiness, images were pooled for Light and Dark *Ai34* line *cFos-tTA^-/-^TetO-Cre^+/-^*mice, and averaged images were subtracted from each corresponding *Ai34* line *cFos-tTA^+/-^TetO-Cre^+/-^*image. Z-scoring was applied separately to each resulting image using a warped tissue mask and a hemispheric mask: ([image − mean intensity in the brain]/standard deviation of intensity in the brain). The hemispheric mask spatially constrained voxel-wise analyses by excluding non-somal regions (ventricles, fiber tracts), undefined edge voxels, and hindbrain/brainstem regions with poor *Ai14* line transcriptional induction in the cerebellum, pons, and medulla.

*Ai14* line tdTomato^+^ somata were segmented with Ilastik ^62^ (v1.4.0; pixel classification) for brain-wide cell counts using connected-components-3d (v3.16.0; connectivity = 6), yielding centroid coordinates for each cell. Cell centroids were converted to 50µm images for binning detections. Images were warped to 30µm atlas space, smoothed, and hemispheres were averaged together. To correct for background, images were pooled for Light and Dark *Ai14* line *cFos-tTA^-/-^TetO-Cre^+/-^*mice, cell counted images were averaged across hemispheres and pooled. The averaged cell count images were subtracted from each corresponding *Ai14* line *cFos-tTA^+/-^TetO-Cre^+/-^*sample. *Ai14* line cell densities were then z-scored across each brain.

Voxel-wise differences in mean signal between the Light and Dark groups were assessed using a two-sample t-test within a general linear model, with non-parametric permutation inference (18,000 permutations) implemented with FSL’s randomise_parallel (FMRIB; v6.0.2). Voxel-wise differences in variability between the Light and Dark groups were assessed by transforming each subject’s image into absolute deviations from its group’s voxel-wise median. These transformed images were then compared using the same two-sample t-test design and nonparametric permutation procedure used for mean comparisons. For voxel-wise mean and variance difference tests, multiple comparison correction was performed using the false discovery rate method to identify clusters of significant voxels at q < 0.05, q < 0.10, q < 0.15, and q < 0.20 (See Data S1 for all p-value/q-value relationships). Clusters smaller than 100 contiguous voxels were excluded from further analysis to minimize false positives (See Supplemental Fig. 1 for visualization of the voxel-wise analysis pipeline). Statistical maps were saved in the compressed NifTI format (.nii.gz) and visualized in FSLeyes, then converted to .parquet files for analysis, including aggregation of average Cohen’s *d*, average log_2 s_tandard deviation ratio, and percent of significant voxels by anatomical brain region at different levels of subdivision. n = 5 Light and 5 Dark *cFos-tTA^+/-^TetO-Cre^+/-^*and 3 Light and 3 Dark *cFos-tTA^-/-^TetO-Cre^+/-^*pooled background subtraction mice for each reporter line were used for mean and variance assessments.

For assessment of stereotypy under Light conditions, voxel-wise comparisons of standard deviation were performed with a Pitman-Morgan test for difference in variance between dependent groups ^63,64^ between the z-scored, background subtracted *cFos-tTA^+/-^TetO-Cre^+/-^Ai14* and *Ai34* Light group vs. their own voxel-wise spatially scrambled maps, with each map scrambled independently. Scrambled > Original and Original > Scrambled voxels surviving false discovery rate thresholds of q < 0.05, q < 0.10, q < 0.15, and q < 0.20 were mapped, and clusters smaller than 100 contiguous voxels were excluded from further analysis to minimize false positives. n = 5 Light *cFos-tTA^+/-^TetO-Cre^+/-^*and 3 Light and 3 Dark *cFos-tTA^-/-^TetO-Cre^+/-^*pooled background subtraction mice for each reporter line were used for stereotypy assessments.

All Python code used to manipulate .parquet files and to generate tables and quadrant plots can be found at https://github.com/ojlesperance/ai14-ai34-light-dark-whole-brain, and source files can be found at https://zenodo.org/records/23091056 (DOI: 10.5281/zenodo.23091056). Correlation p and *ρ* values were calculated in GraphPad Prism v11.1.0. Scripts from UNRAVEL, used for preprocessing and analysis, are available at zenodo.org/records/13655988, but see github.com/b-heifets/UNRAVEL for the latest code. Interactive versions of statistical maps shown in Figs. 2,3, and 5 can be found at https://heifetslab-unravel.org/.

**Figure 1:**
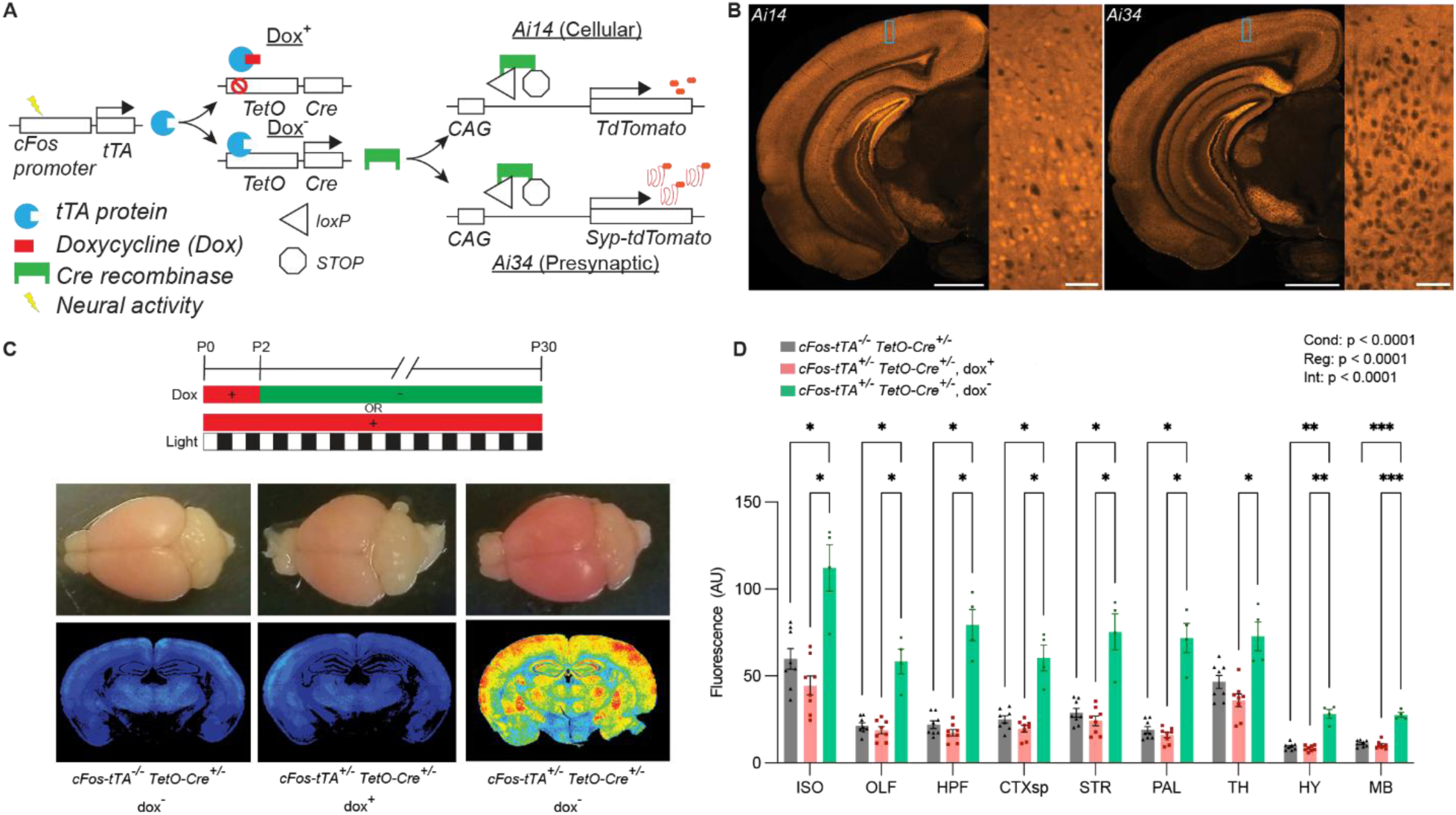
Signal and background characterization of triple transgenic somal and presynaptic labeling strategies. **A)** Schematic of temporally controlled, *cFos* promotor-dependent pharmacogenetic strategy for labeling cell bodies (*Ai14*) or presynaptic terminals (*Ai34*). **B)** Representative whole hemisphere (left panel) and visual cortical (right panel) images of labeled cell bodies (*Ai14*, left) or presynaptic terminals (*Ai34*, right) of triple transgenic mice housed under a standard 12h light/dark cycle. Scale bar: 1mm (left panel), 100µm (right panel). **C)** Postnatal to P30 recording timeline (above), top-down whole-brain images (middle), and 16-color heatmaps of tdTomato fluorescence from a coronal slice (below) of *cFos-tTA^-/-^TetO-Cre^+/-^*, cFos*-tTA^+/-^TetO-Cre^+/-^*dox^+^, and *cFos-tTA^+/-^, TetO-Cre*^+/-^dox^-^presynaptic line brains. **D)** Quantification of fluorescence in major brain area subdivisions of *cFos-tTA^-/-^TetO-Cre^+/-^*, *cFos-tTA^+/-^TetO-Cre^+/-^*dox^+^, and *cFos-tTA^+/-^TetO-Cre^+/-^*dox^-^presynaptic line mice. *p<0.05, **p<0.01, ***p<0.001, 2-way ANOVA effects of experimental condition (Cond.), brain region (Reg.), interaction (Int.) and Tukey’s multiple comparisons results for each group are shown. n=8 *cFos-tTA^-/-^TetO-Cre^+/-^*, 8 *cFos-tTA^+/-^TetO-Cre^+/-^*dox^+^, and 4 *cFos-tTA^+/-^TetO-Cre^+/-^*dox^-^mice.

**Figure 2:**
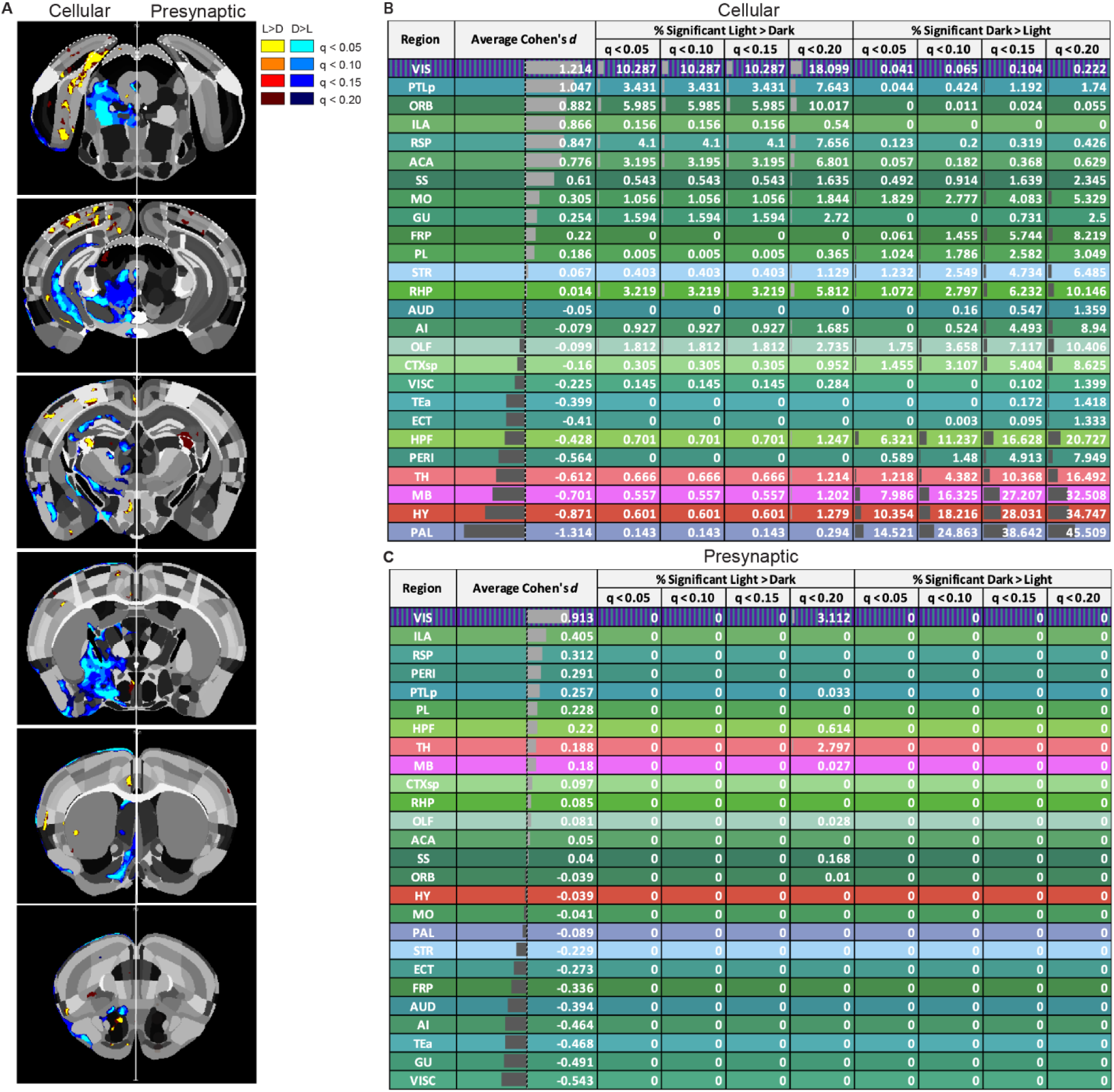
Early visual deprivation triggers subcortical-biased cellular *cFos* expression. **A)** Cellular (left) and presynaptic (right) distributions of voxels with significant differences in background subtracted and z-scored *cFos*-driven tdTomato expression between Light (12h light/dark cycle) and Dark (24h visual deprivation) conditions. The visual cortex, lateral geniculate nucleus, and sensory superior colliculus are indicated with dotted lines. Voxels reaching FDR-corrected thresholds of q < 0.05, q < 0.10, q < 0.15, and q < 0.2 are shown. Average Cohen’s *d* (Light-Dark) for all voxels in the indicated region and the percentage of region volume containing statistically significant voxels at each FDR threshold are shown for **B)** cellular and **C)** presynaptic line mice. n = 5 presynaptic and 5 cellular line *cFos-tTA^+/-^TetO-Cre^+/-^*mice each for Light and Dark conditions. n = 6 presynaptic and 6 cellular line *cFos-tTA^-/-^TetO-Cre^+/-^*mice used for background subtraction. Voxel-wise permutation-based mean difference test with FDR correction, 18,000 permutations.

**Figure 3:**
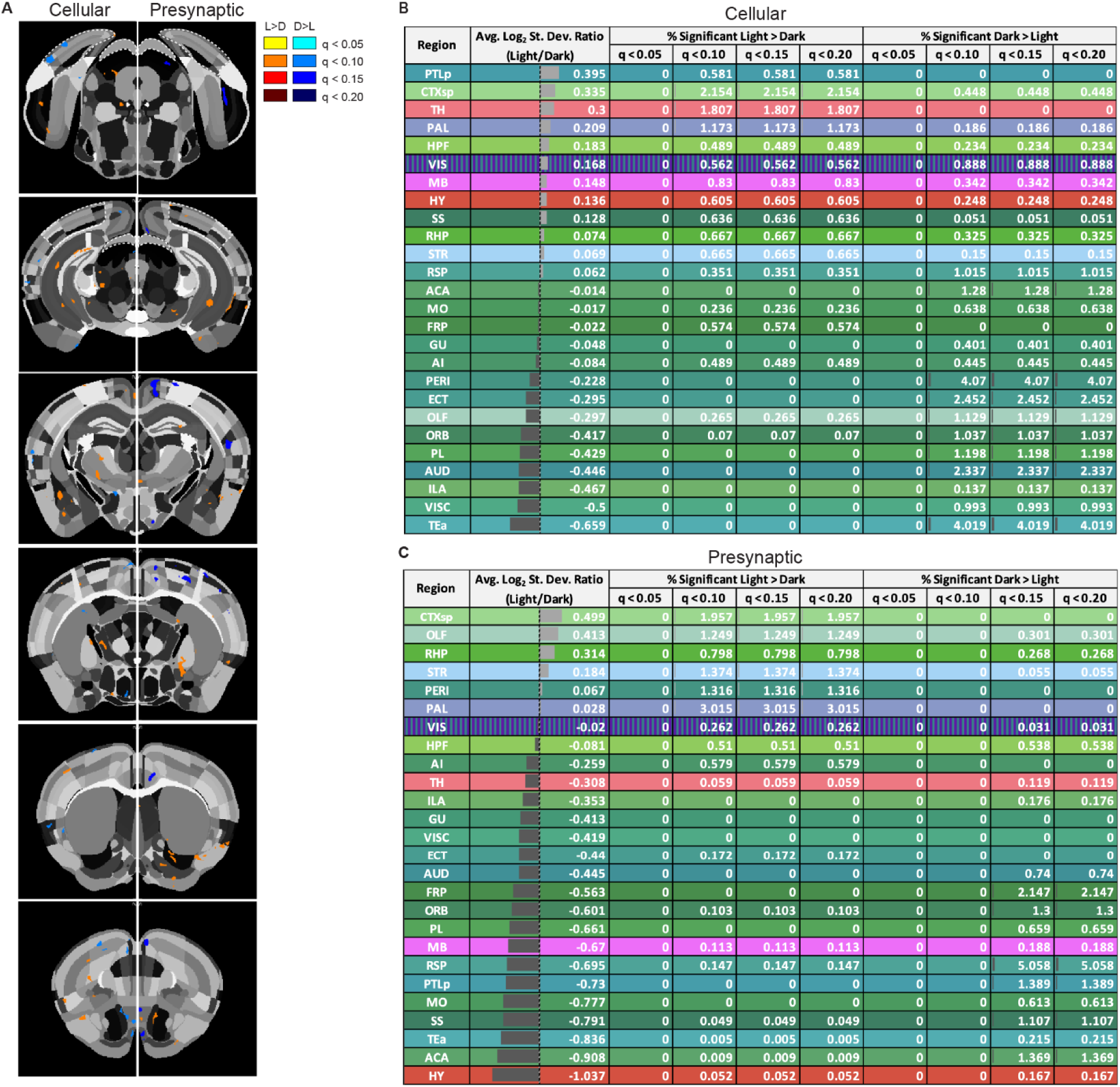
Early visual deprivation weakly alters between-subject variance. **A)** Cellular (left) and presynaptic (right) distributions of voxels with significant differences in between-subject variance of background-subtracted, z-scored *Ai14* line tdTomato^+^ cellular density or *Ai34* signal, respectively, between Light (12h light/dark cycle) and Dark (24h visual deprivation) conditions. The visual cortex, lateral geniculate nucleus, and sensory superior colliculus are indicated with dotted lines. Voxels reaching FDR correction thresholds of q < 0.05, q < 0.10, q < 0.15, and q < 0.20 are shown. Average log_2 s_tandard deviation ratios (Light/Dark) across all voxels and the percentage of regional volume occupied by significant voxels at each FDR threshold are shown for **B)** cellular and **C)** presynaptic datasets, aggregated by anatomical region. n = 5 cellular-line and n = 5 presynaptic-line *cFos-tTA^+/-^TetO-Cre^+/-^*mice each for Light and Dark conditions. n = 6 presynaptic and 6 cellular line *cFos-tTA^-/-^TetO-Cre^+/-^*mice were used for background subtraction. Voxel-wise permutation-based Levene’s test with FDR correction, 18,000 permutations.

Parvalbumin immunostaining was performed on a separate set of Light and Dark-exposed mice from the serial two-photon imaged mice, n = 8 Light and 9 Dark mice. Parvalbumin neuron immunostained images were mapped onto the Allen mouse brain atlas as described above. Fluorescence intensity (LoG = 55 filter) and cell size thresholds (lower bound = 8µm; upper bound = 30µm) were used to distinguish parvalbumin-positive cells from background or false positives as previously described^65^. Cells identified as meeting the threshold criteria were manually inspected prior to density quantification for each region. Density per mm^2^ was calculated across 26 regions using NeuroInfo (MBF Bioscience; Williston, VT) ^61^. Differences between Light and Dark group parvalbumin^+^ cell density were assessed using a restricted effects maximum likelihood mixed-effects analysis with Geisser-Greenhouse’s epsilon correction (brain region [within mouse], experimental condition [between mice], interaction, matched by mouse) with post-hoc Šídák’s multiple comparisons tests between conditions within each brain area (1 family, 26 comparisons per family, all ns) using GraphPad Prism v11.1.0.

## Results

### Development and characterization of a *cFos*-driven presynaptic and cellular transgenic labeling system

To develop a permanent fluorescent readout of *cFos-*expressing neurons and their presynaptic projections across the brain, we generated two novel triple transgenic mouse lines: one in which tdTomato accumulates in somata (*Ai14*; cellular line) ^59^ and one in which synaptophysin-tdTomato accumulates in presynaptic terminals (*Ai34*; presynaptic line) ^60^ of cells with *cFos* promoter activity (Fig. 1A-B). In individual cells of both mouse lines, *cFos* promoter activity drives tetracycline-controlled transactivator (tTA) protein expression ^39,66^. To temporally control downstream fluorophore expression, doxycycline, introduced through mouse chow (40mg/kg), binds to tTA and prevents transactivation of the system’s second transgene, *Cre recombinase* driven by the *TetO* promoter ^58^. In the absence of doxycycline, tTA activation of the *TetO* promoter drives expression of Cre recombinase protein. Cre excises a *loxP*-flanked STOP sequence on the third fluorophore-producing transgene (*tdTomato* [*Ai14*] or *synaptophysin-tdTomato* [*Ai34*]), allowing for constitutive production of fluorophore in individual cells with *cFos* promoter activity.

To determine whether sufficient signal was detectable over background synaptophysin-tdTomato expression in the first postnatal month, we maintained *Ai34* line *cFos-tTA^+/-^TetO-Cre^+/-^*mice (signal) and their *cFos-tTA^-/-^TetO-Cre^+/-^*littermates (background) on 40mg/kg doxycycline mouse chow from before birth until postnatal day 2 (P2), then provided doxycycline-free mouse chow under a 12h light/dark cycle until transcardial perfusion and brain collection on P30. To additionally determine whether 40mg/kg doxycycline chow was sufficient to prevent synaptophysin-tdTomato expression beyond background expression, we maintained *Ai34* line *cFos-tTA^+/-^TetO-Cre^+/-^*mice on doxycycline chow until P30 (signal “off”; Fig. 1C).

Quantification of synaptophysin-tdTomato fluorescence in 9 major anatomical subdivisions of the brain (Fig. 1D; see Table 1 for brain region abbreviations) showed ∼2-4.5-fold increases in *cFos-tTA^+/-^TetO-Cre^+/-^*, dox^-^signal over cFos-tTA^+/-^TetO-Cre^+/-^, dox^+^ and *cFos-tTA^-/-^TetO-Cre^+/-^* groups (signal “off” and background respectively) and no additional fluorophore expression in *cFos-tTA^+/-^ TetO-Cre^+/-^*, dox^+^ over *cFos-tTA^-/-^ TetO-Cre^+/-^* mice in any brain region.

**Table 1:** Major anatomical brain regions and abbreviations for their subregions, aggregated and color coded based on the 2020 Allen Mouse Brain Atlas Common Coordinate Framework, version 3 (CCFv3). Subregions involved in the initial visual processing stream (visual system; VS) are denoted in purple stripes over their anatomical color coding. Cortical areas are further divided into layers 1, 2/3, 4, 5, 6a, and 6b where applicable. Pons, medulla, cerebellum, fiber tracts, and aqueducts were not included in analyses due to low or absent *cFos* expression.

| Region | Abbrev | Subregion |
|---|---|---|
| Isocortex | ACAd | Anterior cingulate area, dorsal part |
|  | ACAv | Anterior cingulate area, ventral part |
|  | AId | Agranular insular area, dorsal part |
|  | AIp | Agranular insular area, posterior part |
|  | AIv | Agranular insular area, ventral part |
|  | AUDd | Dorsal auditory area |
|  | AUDp | Primary auditory area |
|  | AUDpo | Posterior auditory area |
|  | AUDv | Ventral auditory area |
|  | ECT | Ectorhinal area |
|  | FRP | Frontal pole |
|  | GU | Gustatory areas |
|  | ILA | Infralimbic area |
|  | MOp | Primary motor area |
|  | MOs | Secondary motor area |
|  | ORBl | Orbital area, lateral part |
|  | ORBm | Orbital area, medial part |
|  | ORBvl | Orbital area, ventrolateral part |
|  | PERI | Perirhinal area |
|  | PL | Prelimbic area |
|  | RSPagl | Retrosplenial area, agranular part |
|  | RSPd | Retrosplenial area, dorsal part |
|  | RSPv | Retrosplenial area, ventral part |
|  | SSp-bfd | Primary somatosensory area, barrel field |
|  | SSp-ll | Primary somatosensory area, lower limb |
|  | SSp-m | Primary somatosensory area, mouth |
|  | SSp-n | Primary somatosensory area, nose |
|  | SSp-tr | Primary somatosensory area, trunk |
|  | SSp-ul | Primary somatosensory area, upper limb |
|  | SSp-un | Primary somatosensory area, unassigned |
|  | SSs | Supplemental somatosensory area |
|  | TEa | Temporal association area |
|  | VISC | Visceral area |
|  | VISa | Anterior visual area |
|  | VISal | Anterolateral visual area |
|  | VISam | Anteromedial visual area |
|  | VISl | Lateral visual area |
|  | VISli | Laterointermediate visual area |
|  | VISp | Primary visual area |
|  | VISpl | Posterolateral visual area |
|  | VISpm | Posteromedial visual area |
|  | VISpor | Postrhinal visual area |
|  | VISrl | Rostrolateral visual area |
| HPF (Hippocampal Formation) | APr | Area prostriata |
|  | CA1 | Ammon's horn, field CA1 |
|  | CA2 | Ammon's horn, field CA2 |
|  | CA3 | Ammon's horn, field CA3 |
|  | DG | Dentate gyrus |
|  | ENTl | Entorhinal area, lateral part |
|  | ENTl2 | Entorhinal area, lateral part, layer 2 |
|  | ENTl3 | Entorhinal area, lateral part, layer 3 |
|  | ENTm | Entorhinal area, medial part |
|  | ENTm2 | Entorhinal area, medial part, layer 2 |
|  | ENTm3 | Entorhinal area, medial part, layer 3 |
|  | ENTm6 | Entorhinal area, medial part, layer 6 |
|  | FC | Fasciola cinerea |
|  | HATA | Hippocampo-amygdalar transition area |
|  | HPF | Hippocampal formation |
|  | IG | Induseum griseum |
|  | PAR | Parasubiculum |
|  | POST | Postsubiculum |
|  | PRE | Presubiculum |
|  | ProS | Prosubiculum |
|  | SUB | Subiculum |
| CTXsp (Cortical Subplate) | BLAa | Basolateral amygdalar nucleus, anterior part |
|  | BLAp | Basolateral amygdalar nucleus, posterior part |
|  | BLAv | Basolateral amygdalar nucleus, ventral part |
|  | BMAa | Basomedial amygdalar nucleus, anterior part |
|  | BMAp | Basomedial amygdalar nucleus, posterior part |
|  | CLA | Claustrum |
|  | CTXsp | Cortical subplate |
|  | EPd | Endopiriform nucleus, dorsal part |
|  | EPv | Endopiriform nucleus, ventral part |
|  | LA | Lateral amygdalar nucleus |
|  | PA | Posterior amygdalar nucleus |
| CNU (Cerebral Nuclei) | AAA | Anterior amygdalar area |
|  | ACB | Nucleus accumbens |
|  | BST | Bed nuclei of the stria terminalis |
|  | CEAc | Central amygdalar nucleus, capsular part |
|  | CEAl | Central amygdalar nucleus, lateral part |
|  | CEAm | Central amygdalar nucleus, medial part |
|  | CP | Caudoputamen |
|  | FS | Fundus of striatum |
|  | GPe | Globus pallidus, external segment |
|  | GPi | Globus pallidus, internal segment |
|  | IA | Intercalated amygdalar nucleus |
|  | LSc | Lateral septal nucleus, caudal part |
|  | LSr | Lateral septal nucleus, rostral part |
|  | LSv | Lateral septal nucleus, ventral part |
|  | MEA | Medial amygdalar nucleus |
|  | MS | Medial septal nucleus |
|  | NDB | Nucleus of the diagonal band |
|  | OT | Olfactory tubercle |
|  | PAL | Pallidum |
|  | SF | Septofimbrial nucleus |
|  | SH | Septohippocampal nucleus |
|  | SI | Substantia innominata |
|  | STR | Striatum |
|  | TRS | Triangular nucleus of septum |
| IB (Interbrain) | AD | Anterodorsal nucleus |
|  | ADP | Anterodorsal preoptic nucleus |
|  | AHN | Anterior hypothalamic nucleus |
|  | AMd | Anteromedial nucleus, dorsal part |
|  | AMv | Anteromedial nucleus, ventral part |
|  | ARH | Arcuate hypothalamic nucleus |
|  | ASO | Accessory supraoptic nucleus |
|  | AV | Anteroventral nucleus |
|  | AVP | Anteroventral preoptic nucleus |
|  | AVPV | Anteroventral periventricular nucleus |
|  | BA | Bed nucleus of the accessory olfactory tract |
|  | BAC | Bed nucleus of the anterior commissure |
|  | CL | Central lateral nucleus of the thalamus |
|  | CM | Central medial nucleus of the thalamus |
|  | DMH | Dorsomedial nucleus of the hypothalamus |
|  | Eth | Ethmoidal nucleus of the thalamus |
|  | FF | Fields of Forel |
|  | HY | Hypothalamus |
|  | IAD | Interanterodorsal nucleus of the thalamus |
|  | IAM | Interanteromedial nucleus of the thalamus |
|  | IGL | Intergeniculate leaflet of the lateral geniculate complex |
|  | IMD | Intermediodorsal nucleus of the thalamus |
|  | LD | Lateral dorsal nucleus of thalamus |
|  | LGd | Dorsal part of the lateral geniculate complex |
|  | LGv | Ventral part of the lateral geniculate complex |
|  | LH | Lateral habenula |
|  | LHA | Lateral hypothalamic area |
|  | LP | Lateral posterior nucleus of the thalamus |
|  | LPO | Lateral preoptic area |
|  | MD | Mediodorsal nucleus of thalamus |
|  | ME | Median eminence |
|  | MEPO | Median preoptic nucleus |
|  | MGd | Medial geniculate complex, dorsal part |
|  | MGm | Medial geniculate complex, medial part |
|  | MGv | Medial geniculate complex, ventral part |
|  | MH | Medial habenula |
|  | MMd | Medial mammillary nucleus, dorsal part |
|  | MMl | Medial mammillary nucleus, lateral part |
|  | MMm | Medial mammillary nucleus, medial part |
|  | MMme | Medial mammillary nucleus, median part |
|  | MMp | Medial mammillary nucleus, posterior part |
|  | MPN | Medial preoptic nucleus |
|  | MPO | Medial preoptic area |
|  | MPT | Medial pretectal area |
|  | OV | Olfactory ventricle |
|  | PCN | Paracentral nucleus of thalamus |
|  | PD | Posterodorsal preoptic nucleus |
|  | PF | Parafascicular nucleus |
|  | PH | Posterior hypothalamic nucleus |
|  | PIL | Posterior intralaminar thalamic nucleus |
|  | PMd | Dorsal premammillary nucleus |
|  | PMv | Ventral premammillary nucleus |
|  | PO | Posterior complex of the thalamus |
|  | POL | Posterior limiting nucleus of the thalamus |
|  | PR | Perireunient nucleus |
|  | PRC | Precommissural nucleus |
|  | PS | Parastrial nucleus |
|  | PST | Preparasubthalamic nucleus |
|  | PSTN | Parasubthalamic nucleus |
|  | PT | Parataenial nucleus |
|  | PVH | Paraventricular hypothalamic nucleus |
|  | PVHd | Paraventricular hypothalamic nucleus, descending division |
|  | PVT | Paraventricular nucleus of the thalamus |
|  | PVa | Anterior periventricular nucleus of hypothalamus |
|  | PVi | Intermediate periventricular nucleus of hypothalamus |
|  | PVp | Posterior periventricular nucleus of hypothalamus |
|  | PVpo | Preoptic periventricular nucleus |
|  | Pa | Paraventricular hypothalamic nucleus |
|  | PeF | Perifornical nucleus |
|  | PoT | Posterior triangular thalamic nucleus |
|  | RCH | Retrochiasmatic area |
|  | RE | Nucleus reuniens |
|  | RH | Rhomboid nucleus |
|  | RT | Reticular nucleus of the thalamus |
|  | SBPV | Subparaventricular zone |
|  | SCH | Suprachiasmatic nucleus |
|  | SCO | Subcommissural organ |
|  | SFO | Subfornical organ |
|  | SGN | Suprageniculate nucleus |
|  | SMT | Submedial nucleus of the thalamus |
|  | SO | Supraoptic nucleus |
|  | SPA | Suprafascicular nucleus |
|  | SPFm | Subparafascicular nucleus, magnocellular part |
|  | SPFp | Subparafascicular nucleus, parvicellular part |
|  | STN | Subthalamic nucleus |
|  | SUM | Supramammillary nucleus |
|  | SubG | Subgeniculate nucleus |
|  | TH | Thalamus |
|  | TMd | Tuberomammillary nucleus, dorsal part |
|  | TMv | Tuberomammillary nucleus, ventral part |
|  | TU | Tuberal nucleus |
|  | VAL | Ventral anterior-lateral complex of the thalamus |
|  | VLPO | Ventrolateral preoptic nucleus |
|  | VM | Ventromedial nucleus of the thalamus |
|  | VMH | Ventromedial hypothalamic nucleus |
|  | VMPO | Ventromedial preoptic nucleus |
|  | VPL | Ventral posterolateral nucleus of the thalamus |
|  | VPLpc | Ventral posterolateral nucleus of the thalamus, parvicellular part |
|  | VPM | Ventral posteromedial nucleus of the thalamus |
|  | VPMpc | Ventral posteromedial nucleus of the thalamus, parvicellular part |
|  | Xi | Xiphoid thalamic nucleus |
|  | ZI | Zona incerta |
| MB (Midbrain) | APN | Anterior pretectal nucleus |
|  | AT | Anterior tegmental nucleus |
|  | CLI | Central linear nucleus raphe |
|  | CUN | Cuneiform nucleus |
|  | DR | Dorsal nucleus raphe |
|  | DT | Dorsal terminal nucleus of the accessory optic tract |
|  | EW | Edinger-Westphal nucleus |
|  | ICc | Inferior colliculus, central nucleus |
|  | ICd | Inferior colliculus, dorsal nucleus |
|  | ICe | Inferior colliculus, external nucleus |
|  | IF | Interfascicular nucleus raphe |
|  | III | Oculomotor nucleus |
|  | INC | Interstitial nucleus of Cajal |
|  | IPA | Interpeduncular nucleus, apical subnucleus |
|  | IPC | Interpeduncular nucleus, central subnucleus |
|  | IPDL | Interpeduncular nucleus, dorsolateral subnucleus |
|  | IPDM | Interpeduncular nucleus, dorsomedial subnucleus |
|  | IPI | Interpeduncular nucleus, intermediate subnucleus |
|  | IPL | Interpeduncular nucleus, lateral subnucleus |
|  | IPN | Interpeduncular nucleus |
|  | IPR | Interpeduncular nucleus, rostral subnucleus |
|  | IPRL | Interpeduncular nucleus, rostrolateral subnucleus |
|  | IV | Trochlear nucleus |
|  | IntG | Intermediate gray layer of the superior colliculus |
|  | LM | Lateral mammillary nucleus |
|  | LT | Lateral terminal nucleus of the accessory optic tract |
|  | MA | Medial accessory oculomotor nucleus |
|  | MA3 | Medial accessory oculomotor nucleus 3 |
|  | MB | Midbrain |
|  | MRN | Midbrain reticular nucleus |
|  | MT | Medial terminal nucleus of the accessory optic tract |
|  | NB | Nucleus of the brachium of the inferior colliculus |
|  | ND | Nucleus of Darkschewitsch |
|  | NOT | Nucleus of the optic tract |
|  | NPC | Nucleus of the posterior commissure |
|  | OP | Olivary pretectal nucleus |
|  | PAG | Periaqueductal gray |
|  | PBG | Parabigeminal nucleus |
|  | PN | Paranigral nucleus |
|  | PP | Peripeduncular nucleus |
|  | PPN | Pedunculopontine nucleus |
|  | PPT | Posterior pretectal nucleus |
|  | RL | Rostral linear nucleus raphe |
|  | RN | Red nucleus |
|  | RPF | Retroparafascicular nucleus |
|  | RR | Midbrain reticular nucleus, retrorubral area |
|  | SAG | Sagulum nucleus |
|  | SC | Superior colliculus |
|  | SNc | Substantia nigra, compact part |
|  | SNr | Substantia nigra, reticular part |
|  | Su3 | Supraoculomotor periaqueductal gray |
|  | VTA | Ventral tegmental area |
|  | VTN | Ventral tegmental nucleus |
| OLF (Olfactory) | AON | Anterior olfactory nucleus |
|  | COAa | Cortical amygdalar area, anterior part |
|  | COApl | Cortical amygdalar area, posterior part, lateral zone |
|  | COApm | Cortical amygdalar area, posterior part, medial zone |
|  | DP | Dorsal peduncular area |
|  | NLOT | Nucleus of the lateral olfactory tract |
|  | NLOT2 | Nucleus of the lateral olfactory tract, layer 2 |
|  | NLOT3 | Nucleus of the lateral olfactory tract, layer 3 |
|  | OLF | Olfactory areas |
|  | PAA | Piriform-amygdalar area |
|  | PIR | Piriform area |
|  | TR | Postpiriform transition area |
|  | TTd | Taenia tecta, dorsal part |
|  | TTv | Taenia tecta, ventral part |

### Early visual experience reallocates activity-tagged somal representation from subcortical toward cortical regions

To test how early visual experience-driven activity redistributes *cFos*-tagged neurons (*Ai14*) or their presynaptic terminals (*Ai34*), we housed mice under a normal 12h light/dark cycle from birth until P8, then maintained the mice under 24h visual deprivation from P9-14 to prevent potential effects of visual stimulation through eyelids prior to eye opening. Once all pups underwent initial eye opening by P15, one group of mice was maintained on a 12h light/dark cycle (Light), while another group remained under 24h visual deprivation (Dark) until P28, followed by whole-brain imaging of the postmortem brain using serial two-photon tomography (See Videos 1-2 for visualization of each imaging plane’s signal and autofluorescence, respectively, in a Light group *Ai34 cFos-tTA^+/-^ TetO-Cre^+/-^* brain). We removed doxycycline from both groups four days before eye opening ^39,67^ and maintained them without doxycycline for an additional week (until P21) to tag *cFos* promoter activity during the first week of visual experience. We then added doxycycline back and kept the animals for an additional week (P28) to allow sufficient expression and transport of the cellular or presynaptic reporters.

We first confirmed that the expression of fluorophores in the absence of the *cFos-tTA* transgene did not differ between Light and Dark conditions in the visual cortex, dorsal and ventral parts of the lateral geniculate nucleus of the thalamus, reticular nucleus of the thalamus, or superior colliculus (Supplemental Fig. 1). To quantify redistribution of *cFos*-tagged neurons (*Ai14*) or their presynaptic terminals (*Ai34*), we first performed voxel-wise background subtraction using the corresponding mouse lines lacking the *cFos-tTA* transgene, followed by within-brain z-scoring of tdTomato^+^ cell density (*Ai14* line) or fluorescence (*Ai34* line) signal. In both reporter lines, *cFos-tTA^-/-^TetO-Cre^+/-^*mice, whose reporter signal is independent of *Fos* promoter activity, exhibited proportionally high or low signal across all brain regions, creating strong cross-regional signal correlations across the brain (Supplemental Fig. 2A-B). Therefore, we applied within-animal z-scoring to reduce mouse-to-mouse variability in global reporter levels and facilitate comparison of the relative spatial distribution of *cFos*-tagged neurons or presynaptic labeling across the brain (Supplemental Fig. 2C-D). To identify reliable differences between Light and Dark conditions, we identified clusters of >100 contiguous voxels showing significant differences in mean z-scores between Light vs. Dark conditions across multiple FDR thresholds, with a p-value <0.0007 at the most stringent threshold (Fig. 2A), using a permutation-based voxel-wise difference test ^28^. For the cellular and presynaptic datasets, average Cohen’s *d* (Light – Dark) and the percentages of significant Light- and Dark-biased voxels at each FDR threshold were then aggregated by anatomical brain region (Fig. 2B-C, respectively; see Data S1 for regional summaries across condensed regions, all subregions, all isocortex subregions, and cortical layers; see last sheet of Data S1 for corresponding FDR q-value and uncorrected p-value thresholds).

We found that visual cortical regions (VIS) and upstream regions in the early visual pathway (visual system; VS), including the dorsal lateral geniculate nucleus (LGd), the superior colliculus (SC), and all VIS subregions, showed strong relative enrichment for *cFos*-tagged somata (Cohen’s *d* [Light – Dark] *=* 0.887). At q < 0.05, ∼6% of VS voxels showed greater relative signal in Light than Dark. Several VIS and LGd also showed significant effects across ∼20-25% of their regional volumes (Data S1), exceeding the Light-biased voxel coverage observed in all other major anatomical brain area aggregations. Beyond the visual cortex, posterior cortical areas, including the retrosplenial (RSP) and posterior parietal association (PTLp) cortices, also showed significant relative enrichment (∼4% of their regional volume) of *cFos*-tagged somata (Fig. 2). In contrast, ventral brain areas, including the pallidum (PAL), hypothalamus (HY), midbrain (MB), and the thalamus (TH), exhibited significant enrichment of *cFos*-tagged neurons in the Dark condition. Several MB and HY nuclei showed Dark > Light effects across 60-70% of their regional volume at q < 0.05 (Data S1). In contrast to the redistribution of somata of *cFos*-tagged neurons, the relative distribution of presynaptic terminals remained mostly unchanged between Light and Dark conditions. Despite moderate to strong relative mean shifts based on Cohen’s *d*, particularly in VS subregions (*d* = 0.913), no presynaptic line voxel clusters reached significance at q < 0.15 in either direction. These results indicate that visual deprivation redistributes *cFos*-tagged somata without comparable presynaptic reorganization, potentially because axons from different source populations converge on common target regions.

### Early visual experience does not alter the spatial stereotypy of *cFos*-tagged somata and their presynaptic terminals

The experience-dependent reorganization of cFos-tagged somata could be spatially stereotyped or heterogeneous across subjects at finer scales. Similarly, the relative stability of presynaptic topography could reflect either shared spatial constraints across subjects or heterogeneous individual patterns that converge at the regional level.To test whether visual experience shifts cross-individual variability in the spatial distribution of *cFos*-tag (stereotypy), we applied a permutation-based voxel-wise Levene’s test for homogeneity of variance to z-scored *Ai14* line tdTomato^+^ somal density and Ai34 signal in the same set of Light- and Dark-reared somal and presynaptic cohorts. Voxels showing significant differences in variance were mapped across multiple FDR thresholds with a p-value <0.0007 at the most stringent threshold (Fig. 3A). The average log_2 s_tandard deviation ratio (Light/Dark) and percentages of significant Light-variable or Dark-variable voxels at each FDR threshold were then aggregated by anatomical brain region (Fig. 3B-C respectively; see Data S1 for regional summaries).

We found that even among brain regions showing the largest Light–Dark differences in variance (log_2 s_tandard deviation ratios), only a small fraction of voxels (at q<0.1) exhibited significant experience-dependent changes in between-subject variance. Regional shifts in presynaptic variability spanned a wider range of log_2 s_tandard deviations ratios (Light/Dark; −1.04 to 0.50) than cellular variability shifts (−0.66 to 0.40; Fig. 3). Despite visual deprivation shifting the allocation of *cFos*-tagged somata away from VS regions, these regions exhibited weak shifts in between-subject variability for presynaptic and cellular measures (log_2 s_tandard deviation ratios of −0.24 and 0.23, respectively; Data S1). Though the experience-dependent changes in stereotypy were limited, some regions showed greater modulation than others. For instance, the auditory cortex (AUD), which primarily processes auditory information, and the temporal association cortex (TeA), which processes multiple sensory signals, including auditory signals, exhibited a Light-dependent increase in stereotypy of *cFos*-tagged somata: ∼2-4% of voxels showed greater between-subject variance in Dark compared to Light, whereas no voxel clusters showed greater variance in Light mice at q<0.1 (Fig. 3). In contrast, Light increased the between-subject variance in ∼1-2% of somal voxels within subcortical structures, such as the thalamus (TH) and pallidum (PAL), with the PAL also exhibiting increased Light-dependent variance in ∼3% of presynaptic voxels (Fig. 3). Together, these results show that visual experience alters stereotypy weakly and only within spatially restricted portions of individual brain regions.

Although visual experience weakly perturbs between-subject variance, regions that show greater reorganization may also exhibit greater between-subject variance if this reorganization occurs at different voxel locations across individuals. We plotted Light-dependent changes in mean z-score (indicating within-brain reallocation) against shifts in between-animal variance across 487 subregions for *cFos*-tagged somata and their presynaptic terminals. We examined these relationships across the whole brain, within the isocortex and midbrain (Fig. 4A–C). Across the brain, regions fell into four categories based on the direction of vision-dependent shifts in relative mean and between-subject variance: higher relative mean and greater variance in Light (quadrant 1; Q1), higher relative mean in Dark but greater variance in Light (Q2), higher relative mean and greater variance in Dark (Q3), or higher mean in Light but greater variance in Dark (Q4). Regions across the brain were classified as residing in quadrants 1-4, and their percent residency was calculated for each anatomically based mean/variance relationship plot. These analyses revealed distinct patterns of cellular reorganization across the same anatomical groupings. Visual experience-dependent reorganization of *cFos*-tagged somata occupied a wider range of subregional Cohen’s *d* values (∼-3.5 to 2.0) and a narrower range of log_2 s_tandard deviation ratio values (∼-1.25 to 1.5; Fig. 4A, Data S1). Roughly equivalent fractions of analyzed regions across the whole brain occupied each quadrant. However, cortical regions were concentrated in Q1 and Q4 (74.77%), indicating a relative mean bias toward Light, while the direction of variance shifts was nearly balanced between greater variance in Light and greater variance in Dark (49.45% vs. 50.56%). In contrast, midbrain regions were concentrated in Q2 and Q3 (62.97%), indicating a relative mean bias toward Dark. Together, these patterns suggest that visual experience shifts the allocation of *cFos*-tagged somata towards cortical regions, whereas visual deprivation favors relatively greater representation in the midbrain.

**Figure 4:**
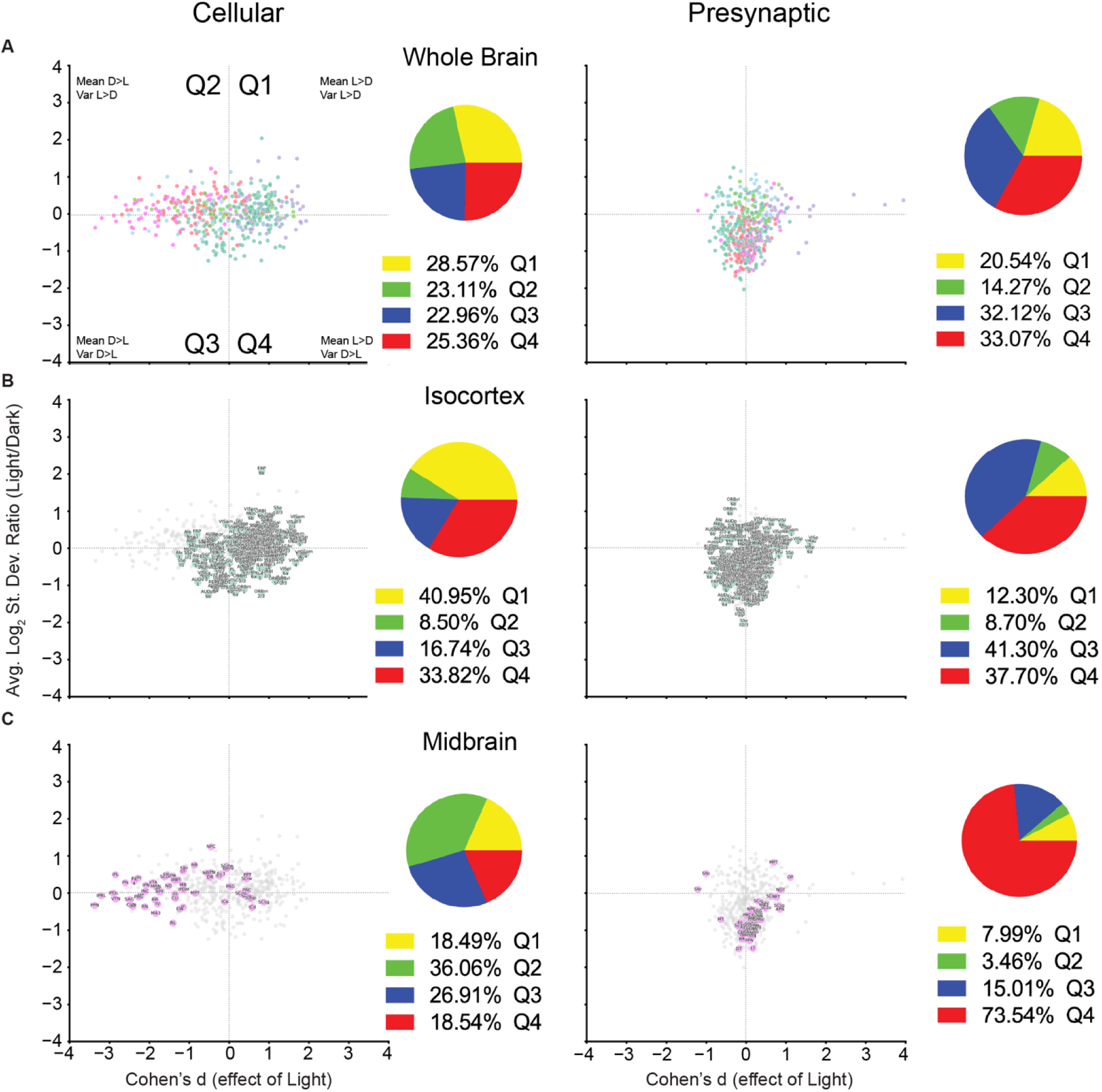
Relationship between experience-dependent shifts in mean spatial allocation and between-subject variance of *cFos*-tagged somata and presynaptic terminals. **A-C** Quadrant plots showing the relationship between Cohen’s *d* (Light - Dark) and the log_2 s_tandard deviation ratio (Light/Dark) across all brain subregions (A), isocortex (B), and midbrain (C) for *cFos-*tagged somata (left) and presynaptic terminals (right). Positive Cohen’s *d* values indicate higher mean z-score in Light, whereas positive log_2 s_tandard devation ratios indicate greater between-subject variance in Light. Anatomical groupings were derived from the Allen Mouse Brain Atlas Common Coordinate Framework (CCFv3; 2020). Colored circles represent brain regions within the indicated anatomical grouping, and the gray circles represent regions outside of it. For each anatomical grouping, pie charts indicate the percentage of voxels residing in each quadrant (color-coded). n = 5 cellular-line and n = 5 presynaptic line *cFos-tTA^+/-^ TetO-Cre^+/-^* mice per Light and Dark conditions. n = 6 cellular-line and n = 6 presynaptic line *cFos-tTA^-/-^ TetO-Cre^+/-^* mice were used for background subtraction.

In contrast to *cFos*-tagged somata, their presynaptic terminals varied predominantly along the variance axis, (∼-2.0 to 1.25 log_2 s_tandard deviation ratio) rather than along the relative-mean-shift axis (∼-1 to 1 Cohen’s *d*; Fig. 4A), with the only exceptions being the three subregions of the LGd (2.70 to 3.95 Cohen’s *d*; Data S1), which receive synaptic input directly from the retina. Presynaptically, most voxels across the brain fell into Q3 and Q4, representing Dark-driven increases in variance (65.19%), with an almost even split between Light and Dark mean biases (Q1 and Q4 vs. Q2 and Q3; 53.61% vs. 46.39%). However, anatomical groupings exhibited specific distributions. The isocortex held the largest proportion of Dark variance-biased presynaptic voxels (79.00 %) with roughly even splits of voxels residing in Q3 and Q4 (41.30% and 37.70%, respectively; Fig. 4B). As expected, the visual system was strongly biased toward higher relative-mean signal (Q1 and Q4) in the Light condition for both somal (Q1 = 50.21%, Q2 = 9.99%, Q3 = 5.72%, Q4 = 34.08%) and presynaptic (Q1 = 33.98%, Q2 = 1.03%, Q3 = 5.40%, Q4 = 59.59%; not shown). Comparison of the cellular and presynaptic brain-wide quadrant plots revealed a shift in the subcortical-cortical gradient from primarily the variance axis presynaptically to the mean axis cellularly, qualitatively resembling a clockwise ∼90° rotation with vertical reflection of the regional distribution.

To test whether dark rearing disrupts parvalbumin (PV) neuron maturation, as has been shown in the visual cortex ^68–71^, we next asked whether regions occupying different quadrants showed consistent changes in PV cell density between Light and Dark mice. A mixed-effects analysis of PV density across 26 representative regions for all quadrants (Supplemental Fig. 3) found no consistent effect of experimental condition or condition × brain region interaction, though a sharp increase in PV density was observed in the medial mammillary nucleus (MM; Q2) in the Dark group, and moderate but variable decreases were observed in some deep layer AUDv and higher-order visual association regions in Q3 and Q1. Mixed-effects analyses binned within each quadrant likewise yielded no significant effect of experimental condition or condition × brain region interaction (not shown).

### Spatial stereotypy exists at cellular scale across the brain

The relatively limited effects of visual deprivation on spatial stereotypy could reflect a weak level of stereotypy already present under normal visual experience, leaving little organization for deprivation to disrupt. To test this, we compared Light-reared mice with independently spatially scrambled versions of the same maps. For cellular and presynaptic lines, we identified clusters of at least 100 voxels with significant variance shifts using a Pitman-Morgan test for dependent samples ^63,64^ across multiple FDR thresholds (Fig. 5A). Average log_2 s_tandard deviation ratios (Scrambled/Original) and the percentage of region covered by significant voxels were binned by anatomical brain region (Fig. 5B-C; Data S1). Positive log_2 r_atios indicate greater variance after scrambling and therefore greater spatial stereotypy in the original Light-reared mice. For *cFos*-tagged somata, ∼30% of voxels in the midbrain and hypothalamus exhibited greater stereotypy than expected after spatial scrambling (q<0.1), whereas, at the same q-threshold, only ∼1% of voxels in these regions showed differences from visual experience. The contrast was stronger for presynaptic terminals: at q < 0.1, ∼65-75% of voxels in these regions exhibited greater stereotypy than scrambled, compared with <3% showing experience-dependent changes in stereotypy. With more stringent criteria (q<0.05), up to 30% of regional and up to 70% of subregional presynaptic line voxels displayed higher stereotypy than scrambled, whereas <2% of somal line regional voxels and <11% of subregional voxels displayed experience-dependent modulation of stereotypy at that level of stringency (Fig. 5B-C, Data S1). Though the majority of the most intrinsically stereotyped subregions resided in thalamic or midbrain regions, several layer 6a and 6b AUD, TEa, and perirhinal cortex (PERI) subregions exhibited extremely high degrees of presynaptic stereotypy (∼60-70% scrambled > original coverage at q < 0.05; Data S1). The VIS showed very weak somal stereotypy (0% at q<0.05 and 2.70% at q<0.1; Fig. 5B). Similarly, the fraction of presynaptic voxels in aggregated VS regions exhibiting greater stereotypy than scrambled was larger (7.89% at q<0.05 and 33.47% at q<0.1) than the fraction showing experience-dependent changes in stereotypy (0% at q<0.05 and 0.15% at q<0.1; Data S1). In contrast, other brain regions, particularly frontal and associational cortices, showed poor spatial stereotypy in baseline conditions, with virtually no voxels showing greater somal stereotypy than expected, and similarly few showing modulation of stereotypy based on visual experience. Together, these results indicate that stereotypy exists at the level of somata and synapses, though it is heterogeneous across the brain.

**Figure 5:**
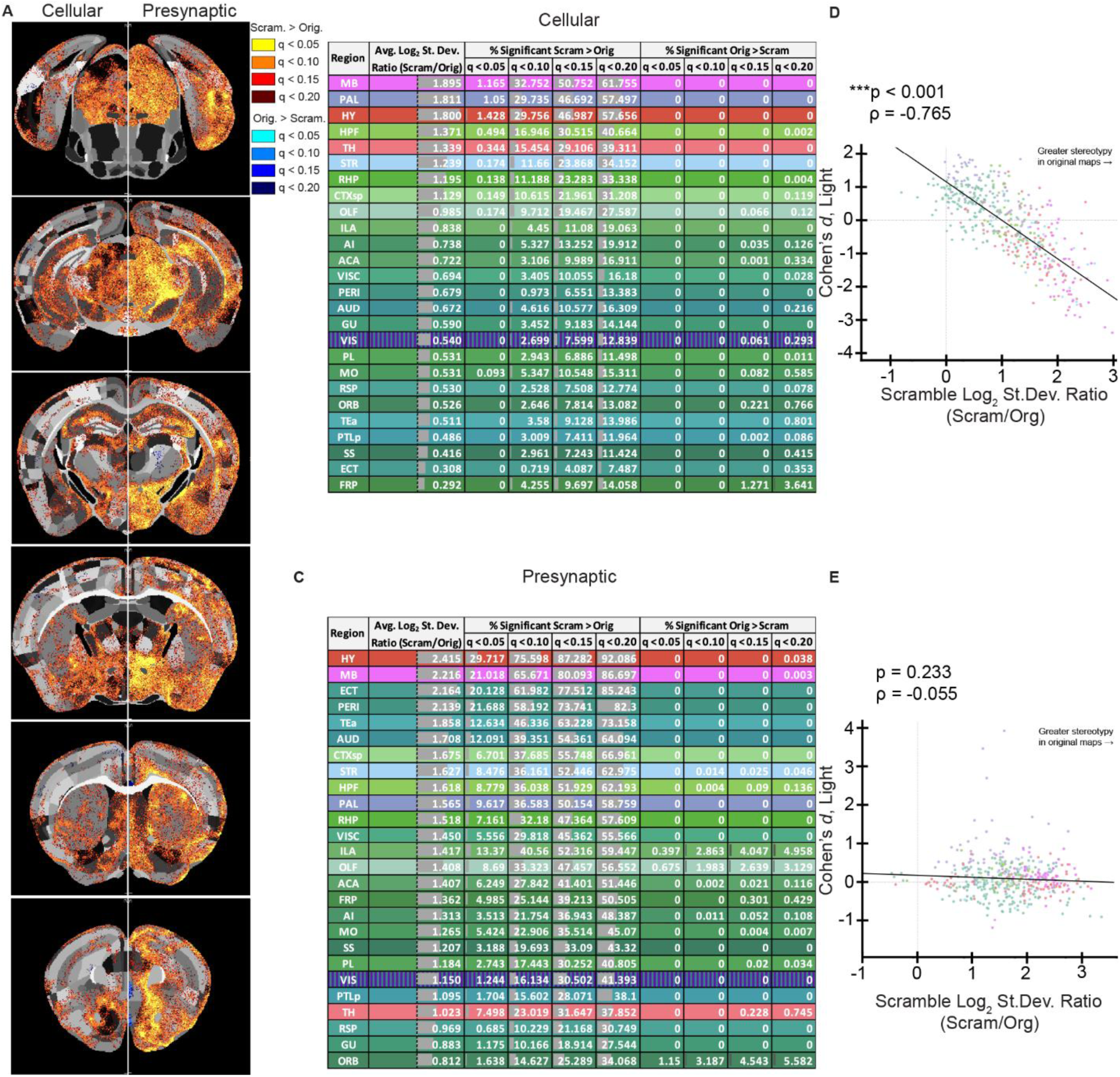
Microscale stereotypy under normal visual experience. **A)** Cellular (left) and presynaptic (right) distributions of voxels with significant differences in variance between original and spatially scrambled images from *cFos-tTA^+/-^ TetO-Cre^+^*^/-^ mice maintained under the standard 12h light/dark cycle (Light condition), following background subtraction and z-scoring of each line. Voxels reaching FDR correction thresholds of q < 0.05, q < 0.10, q < 0.15, and q < 0.20 are shown. Greater variance in scrambled than original maps indicates greater spatial stereotypy in the original maps. Average log_2 s_tandard deviation ratio (Scrambled / Original) across all voxels and percentage of regional volume occupied by significant voxels at each FDR threshold are shown for **B)** cellular and **C)** datasets, aggregated by region. Voxel-wise Pitman-Morgan test for variance of dependent samples. n = 5 cellular-line and n = 5 presynaptic-line *cFos-tTA^+/-^ TetO-Cre^+/-^* mice, with each original image randomly spatially permutated to generate corresponding scrambled images. **(D-E)** Relationships are shown between regional average log_2 s_tandard deviation ratios (Scrambled / Original) and Cohen’s *d* (Light - Dark) for cellular (D) and presynaptic (E) datasets across 487 region subdivisions. p-values and Spearman’s are shown for each comparison. Each data point represents a brain region and is color-coded by anatomical area. Spearman correlation coefficients and p-values are shown.

Furthermore, the weak visual experience-dependent modulation of voxel-level stereotypy does not necessarily reflect weak baseline stereotypy, particularly for the organization of hypothalamic and midbrain presynaptic terminals.

The degree to which the organization of soma or presynaptic terminals is spatially stereotyped may reflect underlying connectivity constraints that could influence their experience-dependent reorganization. Consistently, we observed a striking inverse correlation between baseline stereotypy and light-dependent mean shifts, in which more stereotyped subcortical regions (MB, HY, PAL, STR) experienced the most extreme mean increases in Dark, whereas less stereotyped cortical regions tended to show greater relative mean signal in Light (Fig. 5D; p < 0.001, ρ = −0.765). In contrast, no such relationship was observed for presynaptic terminals (Fig. 5E; p = 0.233, ρ = −0.055).

## Discussion

Brain organization is spatially stereotyped across individuals, with homologous anatomical and functional territories occupying reproducible positions along the anterior-posterior, dorsal-ventral, and medial-lateral axes ^72–74^. However, functional connectivity stereotypy varies by brain region, with multimodal cortices showing greater intersubject variance than unimodal and subcortical regions ^26,75,76^. Thus, functional stereotypy is greater in brain regions with higher spatial constraints on connectivity and lower in cortical regions that underwent disproportionate evolutionary expansions and have longer developmental windows ^76–78^. Here we show that stereotypical functional organization extends to 30×30×100µm voxels across the brain and is also evident at the level of presynaptic terminals. At this resolution, stereotypy follows a similar anatomical gradient, with midbrain and subcortical regions showing greater stereotypy than frontal and other associational cortices. Although both neuronal somata and presynaptic terminals exhibited fine-scale stereotypy, the extent and spatial distribution of stereotypy differed between them. The presynaptic terminals of *cFos*-tagged neurons displayed more voxels with significant scrambled/original log_2 s_tandard deviation ratios than *cFos*-tagged somata at comparable FDR q-value thresholds, while average log_2 s_tandard deviation ratio ranges were comparable between lines. The difference in measurement of relatively low somal and relatively high presynaptic terminal abundance within a single voxel could contribute to their differences in stereotypy. For instance, somata are counted as discrete cells, prior to z-scoring, whereas presynaptic terminal density is measured as continuous fluorescence across many terminals, which could average over many structures and produce spatially smoother voxel value distributions. However, this is unlikely to be the primary factor because each modality was compared to its own scrambled version. Furthermore, there is a larger range of variance (log_2 s_tandard deviation ratio of scrambled/original) in the presynaptic versus somal maps, with some brain regions showing much greater stereotypy than others. Therefore, biological organization also contributes to the differences in stereotypy between presynaptic and somal organization of *cFos*-tagged neurons. For instance, while the exact neuron that becomes *cFos*-tagged may vary within a given region, neighboring neurons in an area with shared function are likely to project to the same anatomically defined target regions. These afferents can contact both *cFos*-expressing and non-expressing downstream neurons, and individual downstream neurons can receive inputs from many different source neurons. Thus, variability observable at the individual soma level could be lost at the presynaptic level through averaging across many converging axons and synapses, producing a more reproducible presynaptic pattern. Nevertheless, these findings suggest that conserved biological constraints shape the cellular-scale functional organization of neurons and their synapses across subjects.

As expected, we observed that one week of visual deprivation strongly decreases relative somal *cFos*-driven tdTomato expression in primary and higher order visual cortices and the LGd, but not the LGv, LP, or PO of the thalamus ^6^. However, this salient experience has only weak effects on stereotypical organization of *cFos-*expressing somata and their presynaptic terminals across the brain. Part of the reason may be that many brain regions have little reproducible spatial organization for experience to disrupt. But even the brain regions that exhibited high levels of stereotypy did not contain a large fraction of voxels showing experience-dependent modulation. This could be because the cellular and projection architecture of many midbrain and subcortical regions is established before eye opening ^74,79,80^. For instance, molecularly distinct hypothalamic nuclei occupy defined spatial domains during embryonic development ^80^. Similarly, retino-collicular and cortico-collicular projections are topographically organized before the onset of vision ^74,79^. Thus, visual deprivation following eye opening is less likely to alter the spatial coordinates of *cFos*-tagged neurons or their projections, and more likely to modify their recruitment. Thus, microscale stereotypy reflects spatial constraints established during brain development and evolution more than an individual’s experience.

The microscale precision of localization of *cFos*-tagged somata in highly stereotyped brain regions could place a constraint on how they can respond to altered experience. Darkness enhanced the relative density of *cFos*-tagged neurons in brain regions with high stereotypy, with a strong inverse correlation between log_2 s_tandard deviation ratio of scrambled/original and Cohen’s *d* of Light-Dark. Because spatial organization is constrained, these regions may respond by recruiting more neurons when experience is drastically altered. This may explain why visual deprivation was expressed along the mean “within-brain” redistribution (Cohen’s d) axis for *cFos*-tagged somata. However, visual deprivation altered presynaptic terminals more along the “between-subject” variance axis. Since presynaptic terminals from different source regions or neurons can buffer the change induced by reorganization of *cFos*-tagged somata, the mean distribution may not change. However, unlike the soma, which persistently occupies a specific coordinate, axons branch or are pruned ^81–83^, providing more opportunities for experience to influence between-subject variability. Conversely, pruning may shunt initially variable presynaptic sprouting patterns initiated by eye opening in total darkness into a more consistent, partially shared synaptophysin-tdTomato signature by P28. Despite organizational axes shifting orthogonally between somal and presynaptic responsiveness to visual experience, they follow a similar subcortical-to-cortical gradient in which darkness increased cellular representation in ventral and subcortical regions, which also showed a greater decrease in stereotypy. Within the circuitry that receives retinal information, the canonical image-forming circuitry ^84,85^ from the lateral geniculate nucleus to the primary visual cortex to higher-order visual cortical areas shows relative enrichment of *cFos*-tagged somata and their presynaptic terminals under normal visual experience. In contrast, subcortical regions such as LGv and IGL, which participate in non-image-forming visual functions ^84,86^, showed increased relative density of *cFos*-tagged somata in Dark, while presynaptic terminals in these regions remained Light-biased. Together, these patterns suggest that altered visual experience is not necessarily expressed as a uniform increase or decrease in activity, but is transformed differently across neuronal compartments and circuits, changing how sensory input is translated into cellular recruitment and presynaptic organization across the brain.

### Limitations

The novel triple transgenic labeling system allows for brain-wide investigation of neuronal circuitry function, but with technical considerations: 1) *cFos*-driven tdTomato signal is cumulative; therefore, we cannot distinguish between different extents of *cFos* promoter activity. Thus, we capture the shift in the density and spatial consistency of the cFos-expressing neurons rather than neuronal activity levels and neuronal firing variance; 2) The presynaptic maps must be interpreted with a biological caveat: while cell bodies do not move or morphologically change between the end of the recording period and the perfusion date (P21-28), their axons and presynaptic terminals can. While the same population of neurons is tagged in cellular versus presynaptic maps, the presynaptic maps represent the P28, potentially post-pruning status of neurons functionally labeled between P15-21; 3) Lastly, the main and accessory olfactory bulbs, cerebellum, and brainstem areas were not assessed due to sparse elicited *cFos* expression, which would artificially compress z-scores across the areas with robust *cFos* expression that this study focused on. Despite these technical considerations, this system was sufficiently sensitive to detect experience-dependent reorganization of *cFos*-tagged neuronal populations and their presynaptic terminals across the brain. Understanding such cellular-scale organization may also provide insight into neurodevelopmental disorders such as autism spectrum disorder, which is associated with altered macroscale functional stereotypy ^87-^_89._

## Author Contributions

OJL and JS designed the experiments. OJL, NP, TM, and EH performed the experiments. DRR and BDH performed voxel-wise analyses. OJL and JS wrote the paper. DRR reviewed and edited the manuscript. JS supervised the research.

## Supporting information

DataS1

## Acknowledgements

This work was supported by the New Faculty General Research Fund from the University of Kansas to JS.

---

**Video 1: Raw Ai34 synaptophysin-tdTomato signal by serial two-photon tomography.** Representative video showing raw synaptophysin-tdTomato serial two-photon-acquired signal in an individual *Ai34 cFos-tTA^+/-^TetO-Cre^+/-^dox^-^*mouse, caudal to rostral.

**Video 2: Raw blue-channel autofluorescence by serial two-photon tomography.** Representative video showing raw serial two-photon-acquired blue channel autofluorescence signal in an individual *Ai34 cFos-tTA^+/-^TetO-Cre^+/-^dox^-^*mouse, caudal to rostral.

## Supplemental Figures

**Supplemental Figure 1:**
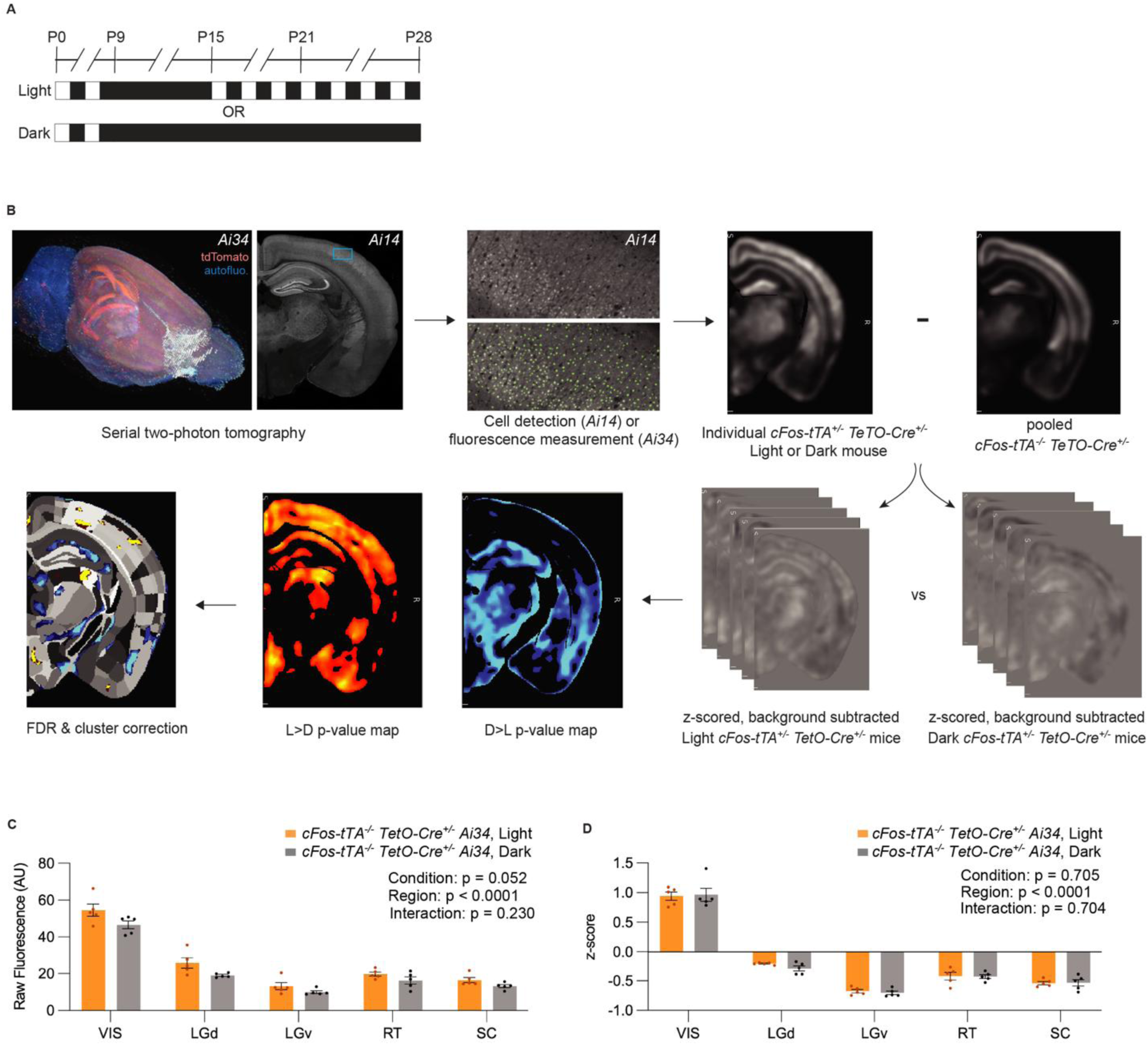
Z-scoring approach and *cFos-tTA^-/-^ TetO-Cre^+/-^* dox^-^ raw and z-scored signal, Light versus Dark. **A)** *cFos*-tag “recording” timeline for Light and Dark group mice. **B)** Representative whole brain (left, *Ai34*) and individual slice (right, *Ai14*) images acquired through serial two-photon tomography. Cell count (*Ai14)* or fluorescence (*Ai34*) is calculated for each 30×30×100μm voxel, then the pooled *cFos-tTA^-/-^ TetO-Cre^+/-^* background map is subtracted from each individual *cFos-tTA^+/-^ TetO-Cre^+/-^* mouse for each respective line. P-value maps are generated by comparing background subtracted, z-scored maps between Light and Dark conditions using UNRAVEL, followed by false discovery rate correction and elimination of clusters of <100 contiguous significant voxels. Raw (**C**) and z-scored (**D**) *Ai34* line *cFos-tTA^-/-^ TetO-Cre^+/-^* signal comparison shows no significant experience-dependent effect of Light vs. Dark exposure in visually responsive areas. n = 5 Light, 5 Dark *Ai34* line *cFos-tTA^-/-^ TetO-Cre^+/-^* mice. 2-way RM ANOVA with Geisser-Greenhouse’s epsilon correction, matched by mouse (Light condition [between mice], brain region [within mouse], interaction) with post-hoc Šídák’s multiple comparisons tests between conditions within each brain area (1 family, 5 comparisons each; all ns).

**Supplemental Figure 2:**
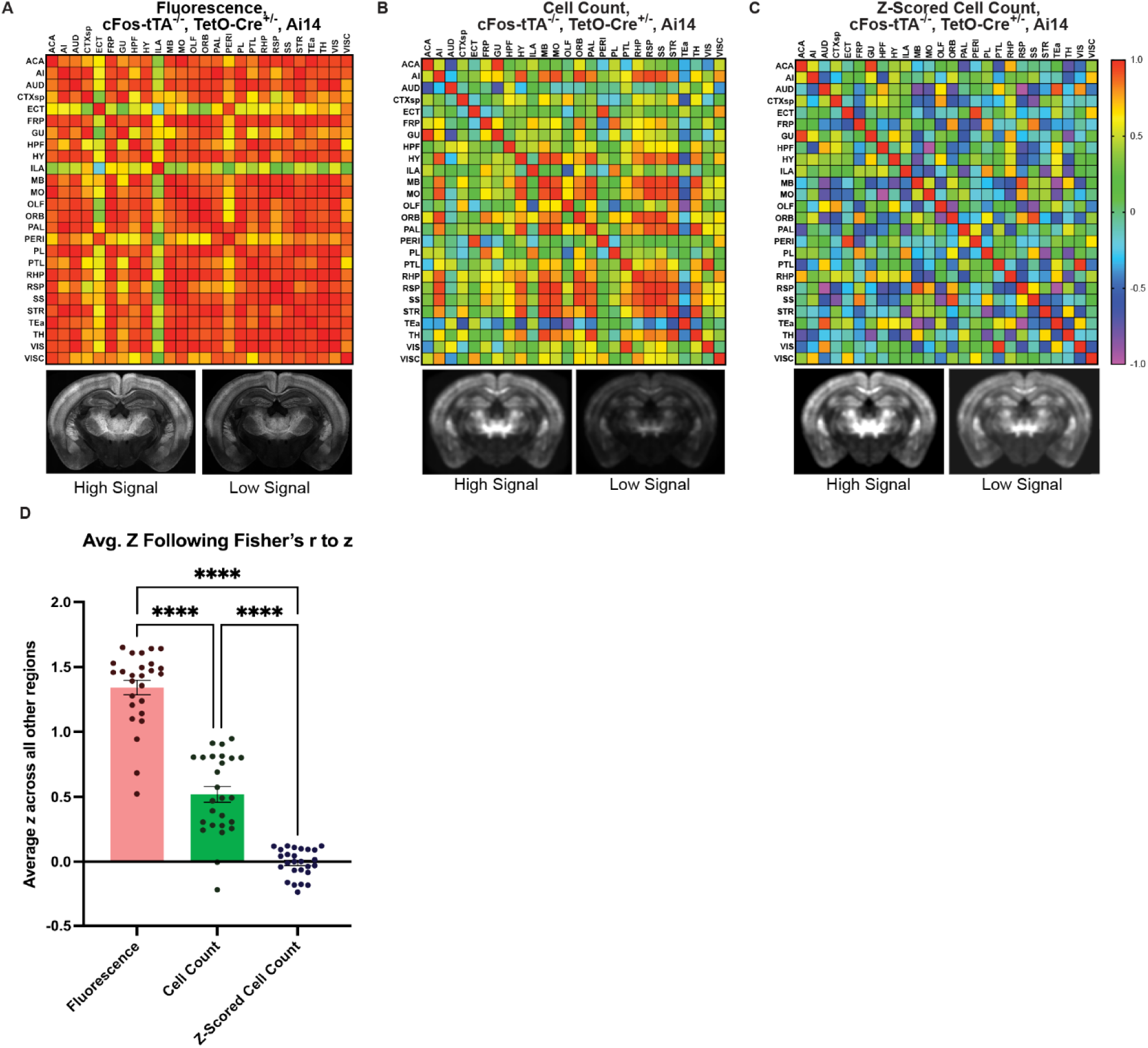
z-scoring *cFos-tTA^-/-^ TetO-Cre^+/-^* Ai14 cell counts reduces the effect of whole-brain mouse-to-mouse TdTomato expression variability in overall fluorescence and cell count. Region-to-region pairwise correlations (Pearson’s *r*; above) in major anatomical brain subdivisions and representative slice images from high and low signal (below) *Ai14 cFos-tTA*^-/-^ *TetO-Cre^+/-^* mice. Images and correlation matrices are derived from **A)** raw fluorescence, **B)** voxel-wise cell count, and **C)** z-scored voxel-wise cell count. Representative images from high and low signal mice are set to identical display ranges for each comparison. **D)** One-way ANOVA (p < 0.0001) with Tukey’s post-hoc multiple comparisons test of average z values from A-C following Fisher’s r-to-z transformation shown. Each data point represents one brain region’s average correlation value with every other region. ****p<0.001. n = 6 *cFos-tTA^-/-^ TetO-Cre^+/-^* cellular line mice (3 Light and 3 Dark, pooled) for all.

**Supplemental Figure 3:**
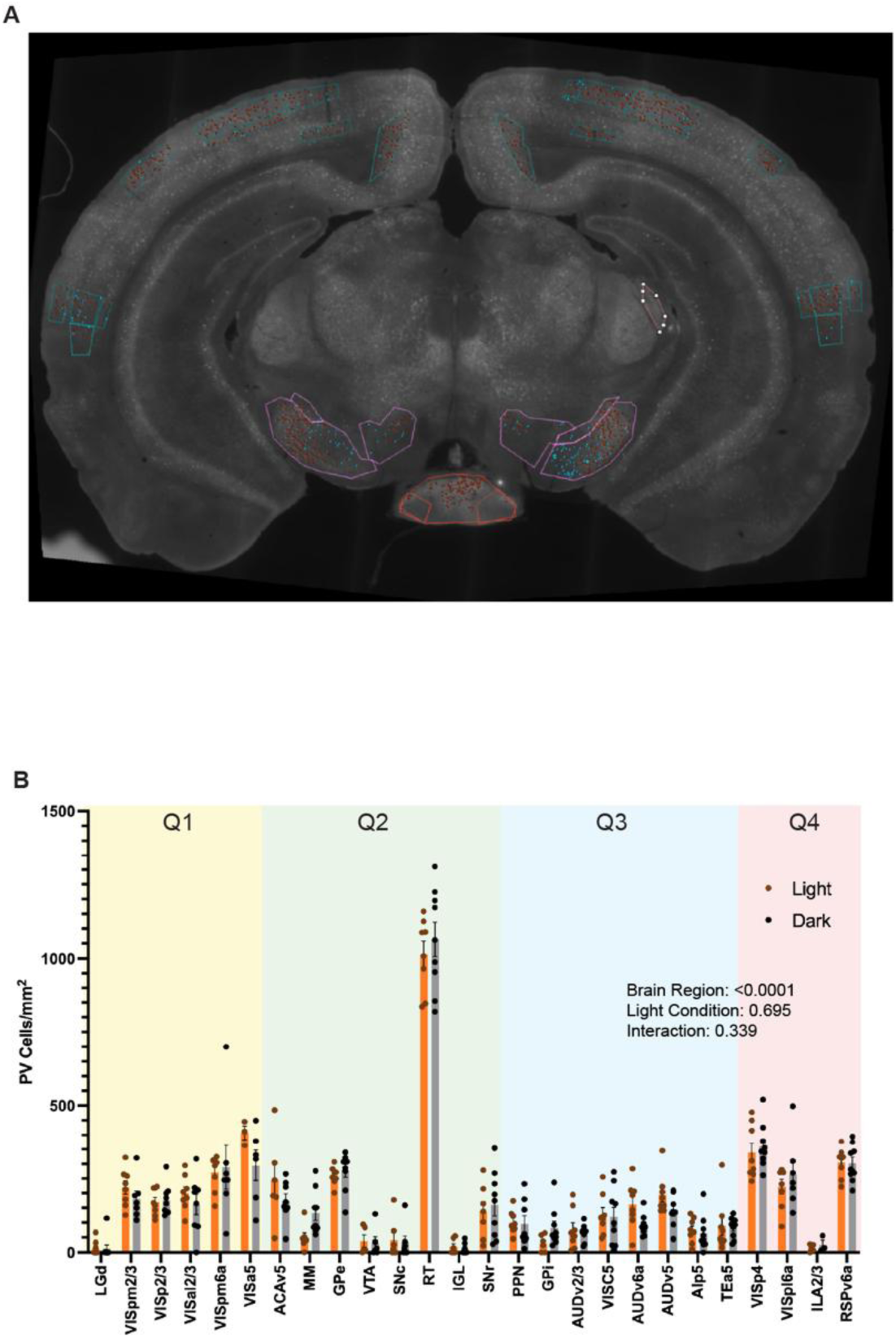
Parvalbumin cell density in regions representing each reorganization quadrant. **A)** Representative slice showing parvalbumin immunohistochemistry and cell detection. Red detection points represent cells algorithmically defined as PV^+^ cells, Blue detection points represent manually defined PV^+^ cells identified during quality control checks. **B)** Raw parvalbumin^+^ cell density across 26 regions representing reorganizational quadrants defined in Fig. 4. n = 8 Light and 9 Dark group mice. Data were analyzed using a restricted effects maximum likelihood mixed-effects model analysis with brain region as a within-mouse factor, experimental condition as a between-mouse factor, and a brain region × condition interaction, with Geisser-Greenhouse correction. *Post hoc* Šídák’s multiple comparisons tests between conditions within each brain area (1 family, 26 comparisons per family, all ns).

**Data S1: Additional regional summaries of voxel-wise analyses and uncorrected p-value thresholds corresponding to FDR q-value thresholds.** Spreadsheets with different regional aggregations (including condensed regions, all subregions, isocortex subregions, and cortical layers) showing average Cohen’s *d* (Light-Dark), average log_2 s_tandard deviation ratio (Light/Dark), average log_2 s_tandard deviation ratio (Scrambled/Original) and the percentage of each region covered by clusters of voxels with significant differences in each of these metrics. The final spreadsheet shows relationship between FDR q-value thresholds and the corresponding uncorrected p-value thresholds for each statistical comparison. n = 5 cellular line *cFos-tTA^+/-^TetO-Cre^+/-^*mice per condition. n = 6 cellular line *cFos-tTA^-/-^TetO-Cre^+/-^*mice for background subtraction. Voxel-wise permutation-based mean difference test with FDR correction, 18,000 permutations for mean and Levene’s tests. For Pitman-Morgan tests, n = 5 original cellular line *cFos-tTA^+/-^TetO-Cre^+/-^*mice, with each image spatially permuted to generate 5 scrambled images. n = 6 cellular line *cFos-tTA^-/-^TetO-Cre^+/-^*mice for background subtraction.

